# Resistant potato starch fuels beneficial host-microbe interactions in the gut

**DOI:** 10.1101/389007

**Authors:** Julian Trachsel, Cassidy Briggs, Nicholas K. Gabler, Heather K. Allen, Crystal L. Loving

**Affiliations:** Food Safety and Enteric Pathogens Research Unit, National Animal Disease Center, Agricultural Research Service, Ames, IA, 50010, USA; Interdepartmental Microbiology Graduate Program, Iowa State University, Ames, IA, 50010, USA; Iowa State University College of Veterinary Medicine Summer Scholar Research Program; Department of Animal Science, Iowa State University, Ames, IA, 50011

**Keywords:** resistant starch, mucosa, microbial accessible carbohydrates, butyrate, microbiota, microbial respiration, prebiotics, t-regulatory cells, colonization resistance, mucosal barrier, immune tolerance

## Abstract

Interactions between diet, the microbiota, and the host set the ecological conditions in the gut and have broad implications for health. Prebiotics are dietary compounds that may shift these conditions towards health by promoting the growth of beneficial microbes. Pigs fed a diet amended with 5% resistant potato starch (RPS) exhibited alterations associated with gut health relative to swine fed an unamended diet (CON). RPS intake increased abundances of anaerobic *Clostridia* in feces and several tissues, as well as intestinal concentrations of butyrate. Functional gene amplicons suggested bacteria similar to *Anaerostipes hadrus* were stimulated by RPS intake. The CON treatment exhibited increased abundances of several genera of *Proteobacteria* (which utilize respiratory metabolisms) in several location. RPS intake increased the abundance of regulatory T cells in the cecum, but not periphery, and cecal immune status alterations were indicative of enhanced mucosal defenses. A network analysis of host and microbial changes in the cecum revealed that regulatory T cells positively correlated with butyrate concentration, luminal IgA concentration, expression of IL-6 and DEF1B, and several mucosa-associated bacterial taxa. Thus, the administration of RPS modulated the microbiota and host response, improved cecal barrier function, promoted immunological tolerance, and reduced the niche for bacterial respiration.

**Importance:** The gut microbiota is central to host health. Many disease states and disorders appear to arise from interactions between the gut microbial community and host immune system. As a result, methods and interventions to support the growth and activity of beneficial gut microbes are being actively pursued. Feeding the gut microbiota with dietary compounds, known as prebiotics, is one of the most promising ways to support gut health. Here we describe a successful prebiotic intervention in weaning swine, a relevant translational model for human health. This work unites microbial and immunological data and demonstrates one way a prebiotic intervention can play out for the benefit of the host.

## Introduction

Dietary prebiotics, such as resistant starches, provide an attractive alternative to antibiotics for improved animal health, including humans (1). Resistant starches are compounds that are not digested by the host in the small intestine, thereby arriving in the large intestine as microbial-accessible carbohydrates. Bacteria that consume resistant starches are normal members of the microbiota and are associated with intestinal health (2). Without access to diet-derived carbohydrates, bacteria will harvest host-derived sugars from the intestinal mucus layer, compromising barrier function (3). However, if dietary carbohydrates are accessible, bacteria will ferment these compounds and release beneficial metabolites, particularly short-chain fatty acids (SCFAs). Host cells consume the vast majority of microbial-produced SCFAs, fueling intestinal homeostasis (4–6).

The SCFA butyrate is a central metabolite for maintaining intestinal homeostasis. It is the preferred fuel source for colonocytes, which oxidize butyrate and consume oxygen. This oxygen consumption lowers the oxygen potential of the epithelia, reducing the amount of electron acceptors available for bacterial respiration (4, 7–9). Furthermore, butyrate can help to limit immune activation by enhancing mucosal barrier function and immunological tolerance, reducing the secretion of immune-derived reactive oxygen and nitrogen species, which also can be used in microbial respiration. Butyrate benefits mucosal barrier function by stimulating increased secretion of mucus, antimicrobial peptides (10), and IgA (6), therefore preventing the translocation of intestinal bacteria that would elicit an immune response. Second, butyrate can induce a more tolerant immune phenotype through the generation of several regulatory immune cell types (11). In total, butyrate-driven changes to the gut microenvironment limit the niche for microbes with respiratory metabolisms (8, 9), allowing microbes that specialize in fermentation to outcompete those that respire, such as *Campylobacter, Salmonella*, and *Escherichia* species (4, 7).

Dietary intake of resistant starches may make the intestinal ecosystem less hospitable for potential pathogens and limit the negative impact of weaning on mammalian health. For example, feeding resistant potato starch (RPS) to nursery-aged piglets enhances some markers of gut health (12). However, the mechanisms by which RPS supports intestinal health in the weaned mammal are poorly defined. The results reported here suggest that dietary RPS encouraged beneficial host-microbe interactions in the hindgut ecosystem. Specifically, RPS stimulated bacterial food webs with functional activities for increased SCFA and butyrate production, which caused subsequent changes in the host intestinal immune status. Immune changes included increased T-regulatory cell types in the cecal tissues, markers of enhanced mucosal barrier function, and reduced amounts of potentially harmful bacteria known to utilize respiratory metabolisms. Collectively, dietary RPS given to pigs at weaning modulated bacterial populations and host immune status to a state associated with enhanced intestinal health.

## Results

### No overt differences in health were observed between the two treatment groups

All pigs appeared healthy throughout the study with no obvious differences in health or behavior observed between the groups. No significant differences in weights were observed either at weaning or necropsy. No gross pathology was observed at necropsy, nor were microscopic pathological changes noted in sections of cecal tissues. These data suggest that all animals in this study were healthy regardless of treatment.

### Bacterial community structure and metabolites differed between the two treatment groups

Both the CON and RPS groups showed similar weaning-related changes in their bacterial communities; however, the communities of the groups became less similar over time. Group fecal bacterial community structures did not differ significantly until day 15, which was associated with a recent dietary phase change (12 days post-weaning), marked by a decrease in the amount of dietary lactose (**Figure 1A, Table S3**). By day 21 the two treatment groups had significantly different fecal bacterial community structures. Group differences in community structures were seen in both the 16S rRNA gene sequence-based analysis, as well as the *but-based* analysis. The size of these effects were driven in part by community dispersion (**Figure 1B**). The 16S rRNA gene-based analysis showed significantly less intra-group variability among the RPS-fed pigs compared to the CON-fed pigs at day 21, and the bacterial community structure of both groups became less dispersed as they matured (**Figure 1B, Table S4**). These results suggest that weaning-associated changes in bacterial communities were more profound than RPS-associated changes, and that RPS intake did not immediately affect the composition of the microbiota, but over time, significantly affected both the overall community composition and the intragroup variability as these communities matured.

**Figure 1:**
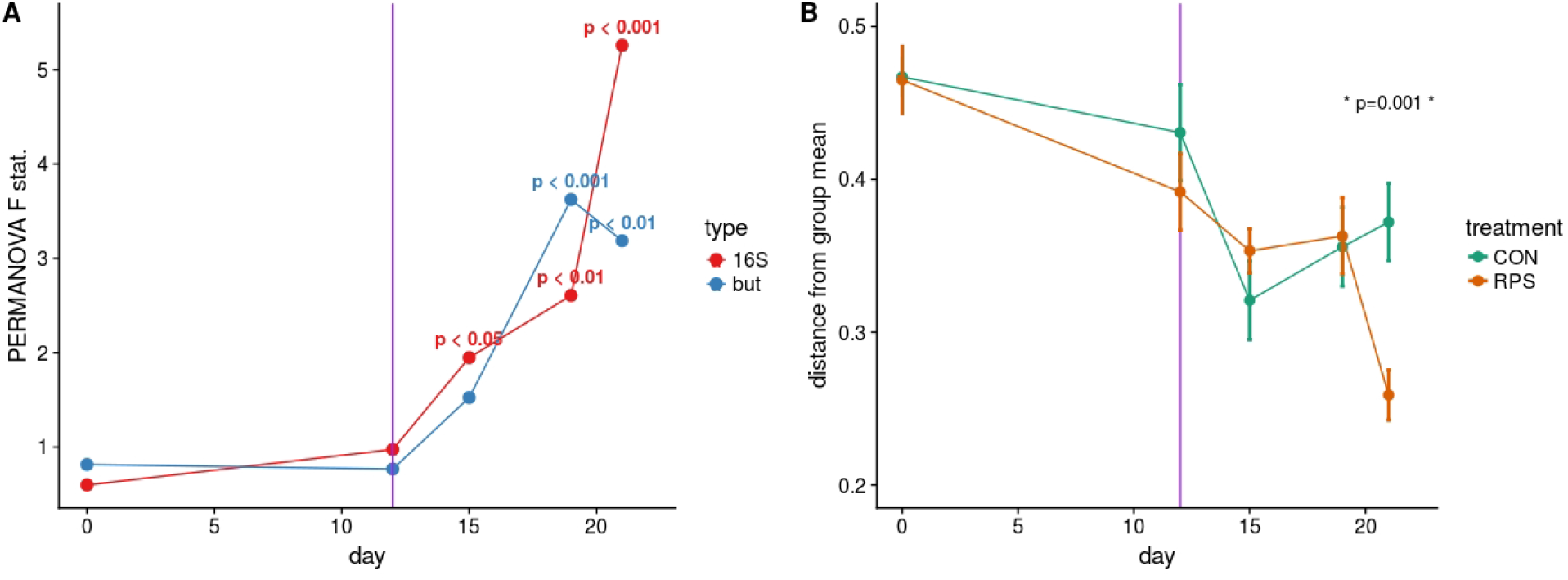
**A:** PERMANOVA pseudo F statistic over time for both the 16S- and *but* gene-based fecal bacterial community analyses. A Bray-Curtis dissimilarity matrix was calculated from OTUs clustered at 97% identity, and this dissimilarity matrix was used as the input for the PERMANOVA tests. Each data point represents the PERMANOVA test statistic (F: intergroup dissimilarity / intragroup dissimilarity) comparing the CON and RPS fecal bacterial community structure at each time point; higher values equate to greater differences between treatment groups. P-values are shown when p < 0.05. **B:** Group dispersion over time. Higher values on the Y axis indicate that communities within that group are more dissimilar from each other. Lower Y axis values indicate that communities in that group are more similar to each other. Error bars represent the standard error around the mean.

Twenty-one days after weaning, the structure of the microbiota was significantly different between the CON and RPS groups at multiple intestinal locations. The 16S rRNA gene-based analyses tended to show greater differences than the *but* gene-based analyses (**Figure 2A, B and Table S3**). The tissue-associated bacterial communities exhibited the same dispersion trend as the fecal communities, with the communities in tissues from RPS-fed pigs exhibiting less group dispersion (**Table S4**). These data suggest that the effects of RPS intake on bacterial communities were not limited to one tissue or location, but bacterial populations throughout the intestinal tract were similarly impacted resulting in higher intragroup similarity in the RPS animals.

**Figure 2:**
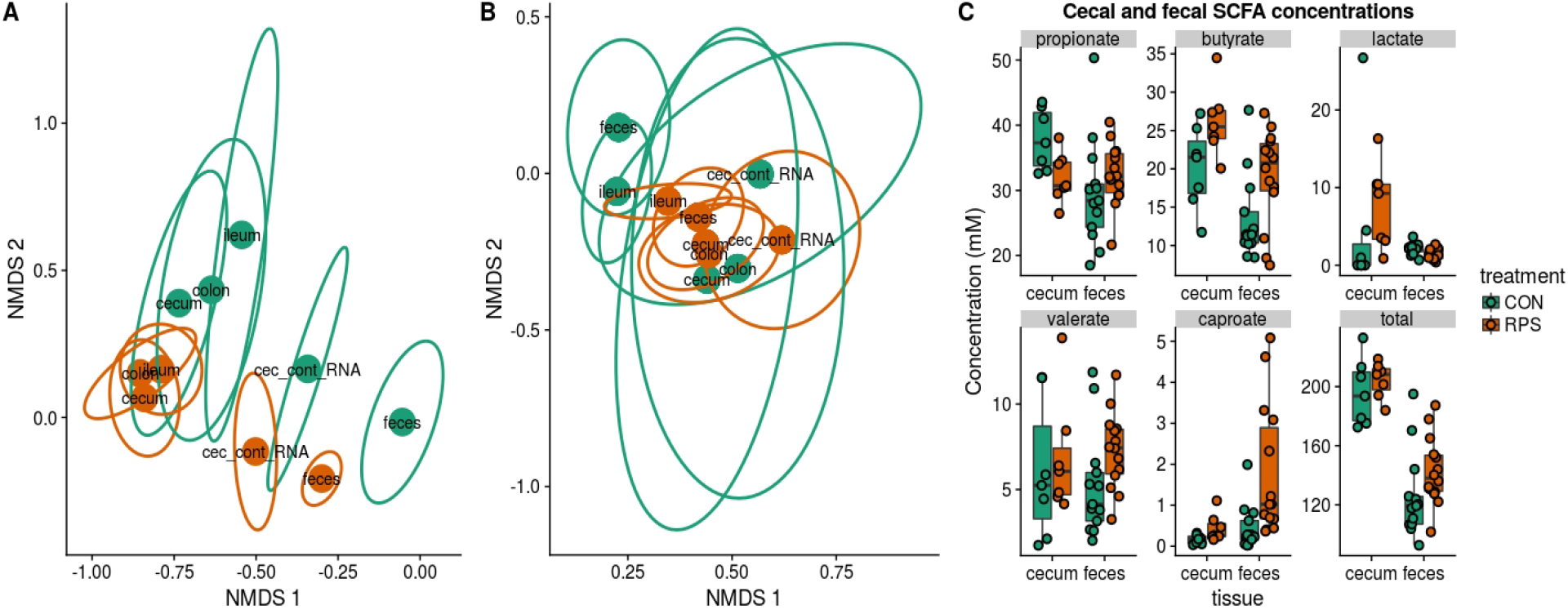
Bacterial community structure and SCFA production in RPS-fed pigs (orange) compared to CON-fed pigs (green) at day 21. Nonmetric multidimensional scaling (NMDS) ordinations of 16S rRNA gene-based Bray-Curtis dissimilarities calculated from OTUs clustered at 97% identity (**A**) and *but* gene-based Bray-Curtis dissimilarities calculated from OTUs clustered at 97% identity (**B**). Points indicate group centroids, from discrete gut locations as labeled. Centroids represent the approximate average community composition, and ellipses show standard error around this average. ‘Cec-cont-RNA’= RNA from cecal contents. **C:** SCFA concentrations (mM) from cecal contents and feces at day 21 post-weaning.

Short-chain fatty acid (SCFA) concentrations in cecal and fecal samples collected at 21-days post-weaning were evaluated to investigate whether the changes in bacterial communities had an impact on microbial metabolic output. Aligning with the changes in microbial community structure, the CON and RPS-fed pigs had differing SCFA profiles (**Fig 2C**). Pigs fed RPS had higher amounts of butyrate in both the cecum and feces (p=0.05 & p=0.05), higher levels of caproate in both the cecum and feces (p=0.07 & p=0.001), lower levels of propionate in the cecum (p=0.05), higher amounts of lactate in the cecum (p=0.09), and lower levels of lactate in the feces (p=0.02). Total SCFA concentrations in cecal contents were not different between the treatments, but total fecal SCFAs were significantly increased in the RPS group (p=0.03). Collectively, dietary RPS modulated bacterial community structure in the distal gut, and these community differences were associated with different SCFA production levels.

### Differentially abundant bacterial genera between the treatment groups

At day 21 post-weaning, many bacterial genera were significantly differentially abundant at several intestinal sites between the treatments (**Figure 3**). Several genera were consistently associated with each respective treatment in most sampling locations. Pigs fed RPS had significantly increased levels of *Terrisporobacter, Sarcina*, and *Clostridium sensu stricto 1* compared to CON-fed pigs. CON-fed pigs had a significant enrichment of *Mucispirillum*, as well as occasional enrichment of various *Proteobacteria* genera such as *Helicobacter, Sutterella*, and *Campylobacter* in different tissues, and many of the genera enriched in CON group harbored members known to utilize respiratory metabolisms (13–16). One 16S rRNA gene OTU, OTU00087, was largely responsible for the differential abundance of the genus *Clostriduim sensu stricto 1* between the two treatment groups. This OTU was greatly enriched in samples from pigs fed RPS, regardless of time point and site (**Figure 4**). Sequences from OTU00087 most closely matched *Clostridium chartatabidum*; organisms similar to this bacterial species are shown to tightly bind starch granules and have been isolated from human feces(17). This suggests that certain bacterial species may occupy the same nutritional niche in many different hosts.

**Figure 3:**
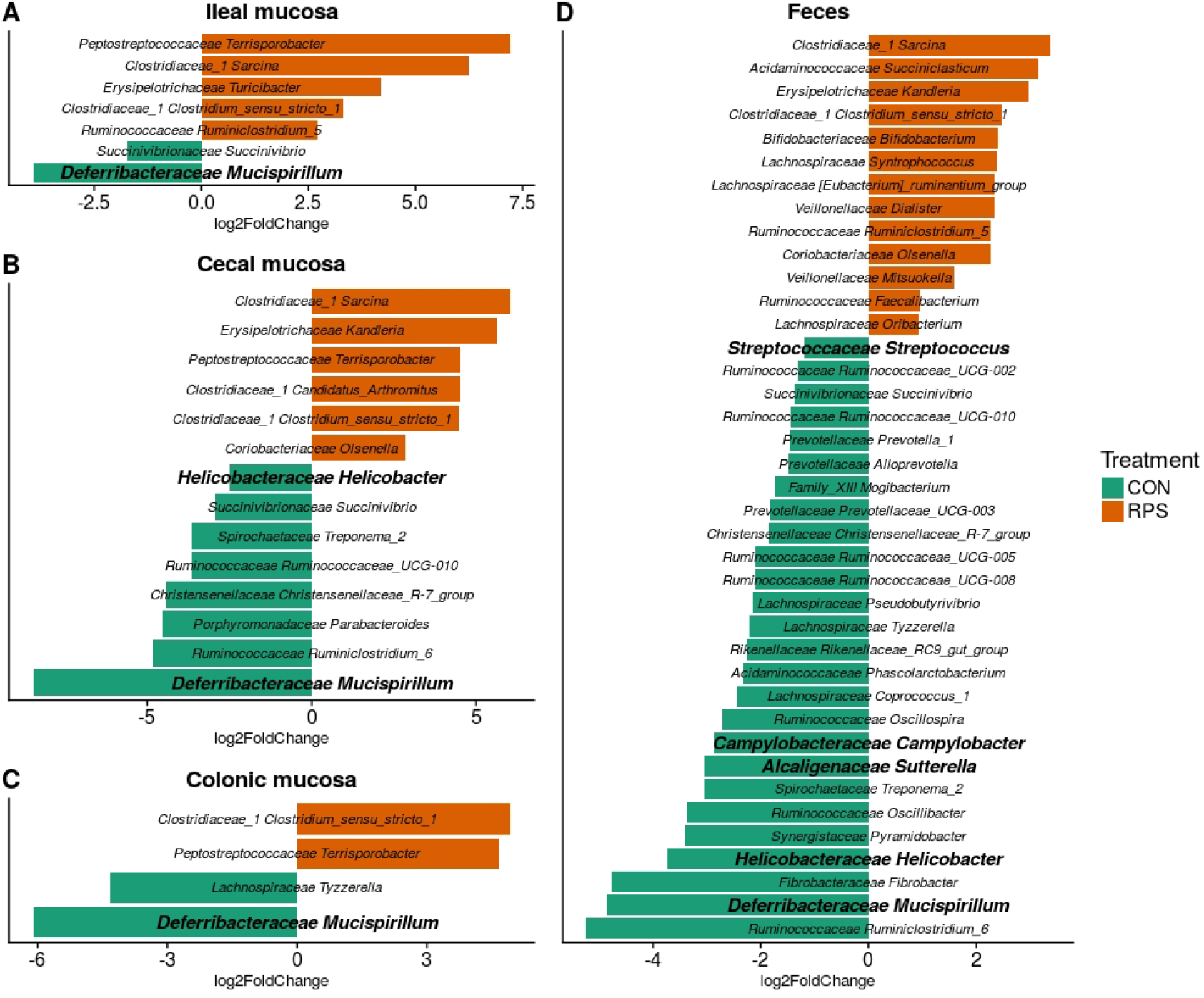
Significantly (p<0.05) Differentially abundant genera based on 16S rRNA gene sequences from the ileal mucosa (**A**), cecal mucosa (**B**), colonic mucosa (**C**), and feces (**D**) as determined by DeSeq2. OTUs clustered at 97% similarity were combined by taxonomic classification at the genus level. The results shown are log2 fold change between the CON (control; green) and RPS (resistant potato starch; orange)-fed groups; note that the x-axis scale is different for each panel. Positive log-fold changes indicate that a genus is enriched in the RPS group, while negative log-fold changes indicate that a genus is enriched in the CON group. The SILVA classification for each genus is labeled on the figure using both the family and genus name. Genera shown in bold are those that harbor species with the capacity for respiration.

**Figure 4:**
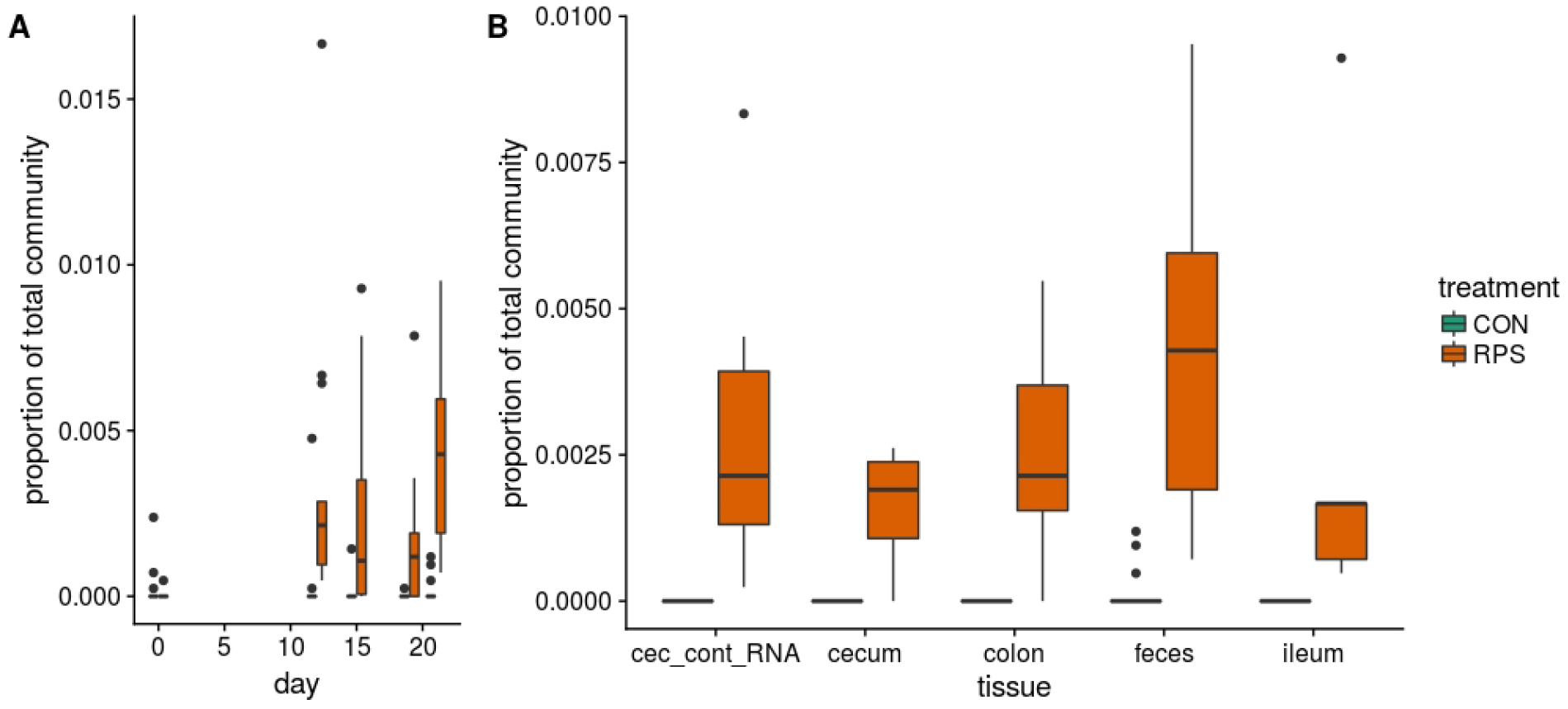
The abundance of OTU0087 over time (**A**) and in intestinal tissues at day 21 (**B**) in CON (green) or RPS (orange)-fed pigs. 16S rRNA gene sequences from this OTU most closely matched those from *Clostridium chartatabidum*, a member of the genus *Clostriduim sensu stricto 1*. Bacteria similar to this organism have been isolated from chemostats fed with resistant starch and inoculated with human feces (17).

Similar to the 16S rRNA gene sequence data, many *but* gene-based OTUs were differentially represented between the two groups (**Figure 5**). RPS intake was consistently associated with a greater abundance of *but* OTUs whose sequences most closely matched to those from *Anaerostipes hadrus*, as well as a *but* OTU whose sequences most closely matched an organism detected in metagenomes from human feces (*but* OTU0067). Intestinal samples from CON group had significant enrichments of *but* OTUs whose sequences most closely matched genes from *Pseudobutyvibrio* and *Helicobacter*. Collectively, these results suggest that particular butyrate-producing bacteria were enriched by RPS intake, and their metabolic activities were likely responsible for increased butyrate concentrations detected in the intestinal samples from these animals.

**Figure 5:**
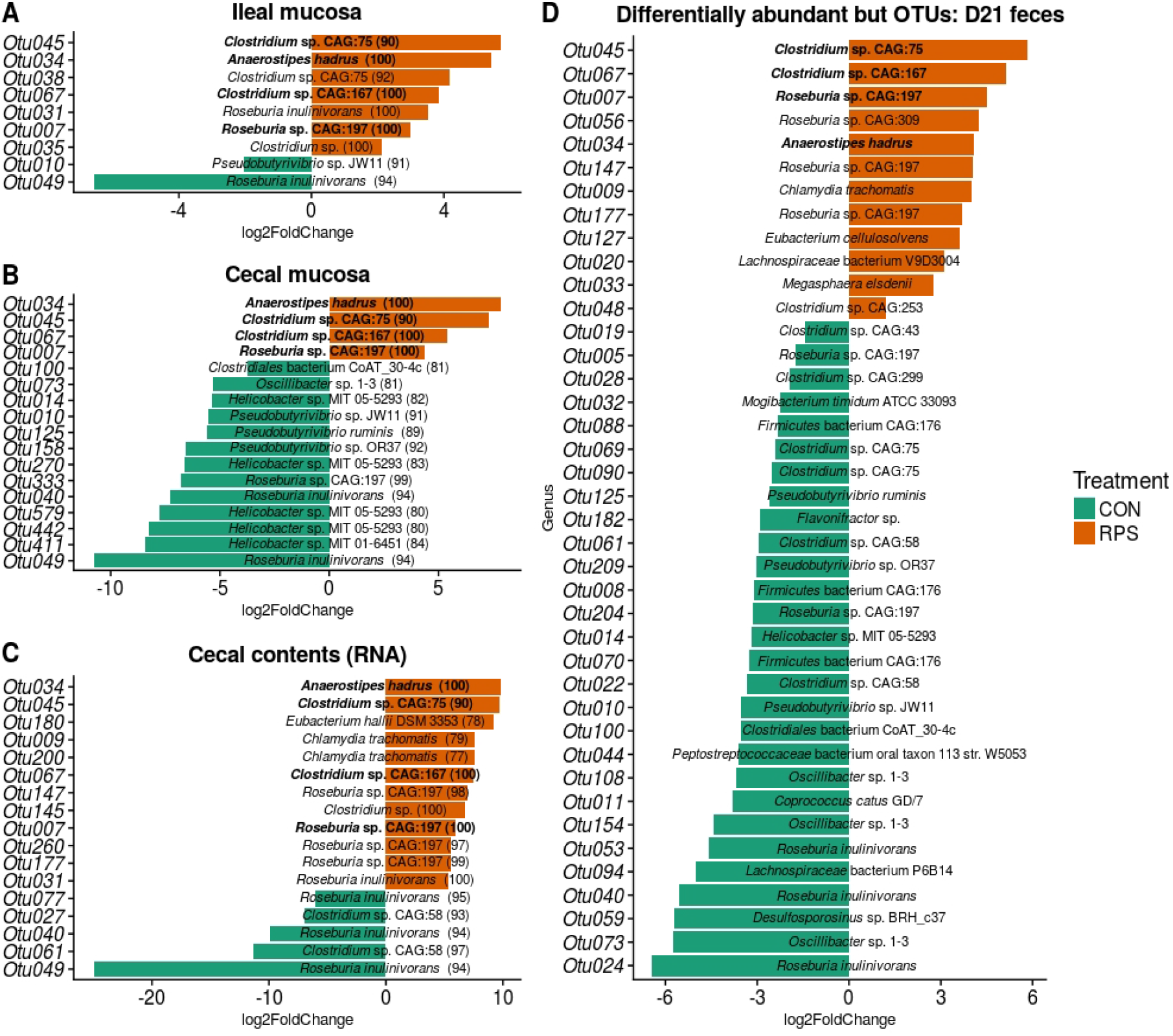
Differentially abundant *but* gene OTUs from the ileal mucosa (**A**), cecal mucosa (**B**), colonic mucosa (**C**), and feces (**D**) as determined by DeSeq2. The results shown are log2-fold change between the CON (control) and RPS (resistant potato starch)-fed pigs; note that the x-axis scale is different for each panel. Positive log-fold changes indicate that an OTU is enriched in the RPS-fed animals, while negative log-fold changes indicate that an OTU is enriched in the CON animals. OTUs were clustered at 97% similarity and are labeled with the species name for their top BLAST hit followed by the percent identity for that hit. OTUs labeled in bold are those that were consistently enriched in one treatment group or the other.

### Bacterial food webs associated with the dietary treatments

To investigate bacterial food webs associated with the increased SCFA levels in the RPS-fed animals, we constructed a correlation network using *but* and 16S rRNA gene OTUs. The network was filtered such that all nodes in the network were differentially abundant between the two groups as determined by DeSeq2, and so the network represented the food webs that were most different between the treatment groups. Nodes enriched in the RPS treatment group clustered into one primary subnetwork, whereas nodes enriched in the CON group clustered into two smaller subnetworks (**Figure 6**). The primary subnetwork enriched in RPS-fed pigs represented bacterial OTUs likely involved in the fermentation of RPS to SCFAs. Bacterial species comprising the RPS-associated network are known to be involved in the breakdown and fermentation of dietary starches, such as *Eubacterium rectale, Mitsuokella* spp., *Prevotella* spp., and *Bifidobacterium* spp (18). These organisms can degrade RPS into small polysaccharides or other metabolic inputs, such as lactate, which are then an available carbon source for other bacteria. Lactate-consuming and butyrate-producing bacteria were also present in the RPS-associated network, including *Megasphaera elsdenii*. These results suggest that dietary RPS supplementation enriched for a bacterial food web that degraded dietary RPS and produced SCFAs via bacterial fermentation.

**Figure 6:**
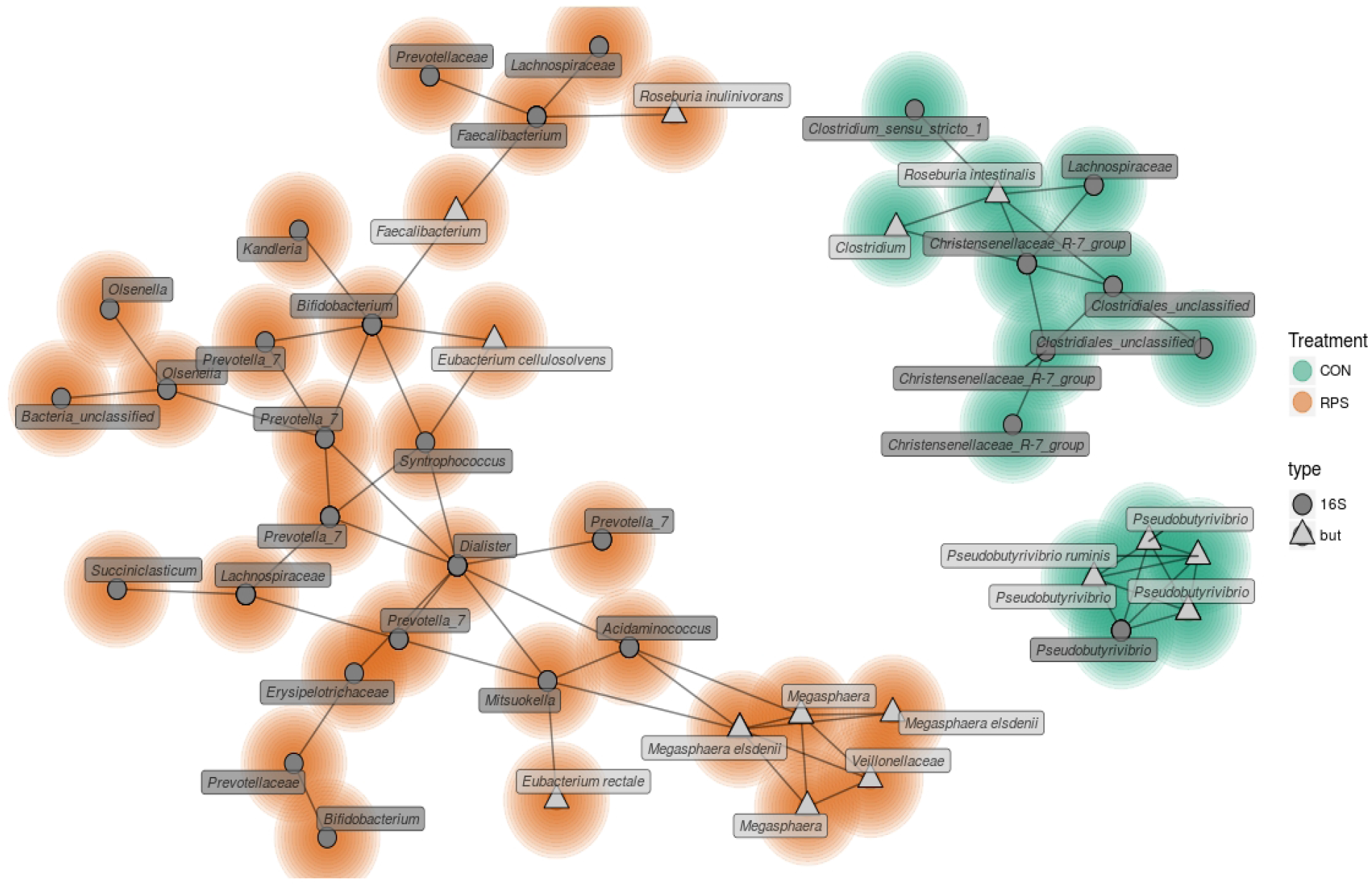
A network depicting correlations between 16S rRNA gene and *but* gene OTUs in the fecal microbiota from RPS and CON fed animals calculated with CCREPE. Only significant (p < 0.05) and positive correlations with a spearman correlation coefficient of at least 0.6 are shown. The network was filtered to only include OTUs that were enriched in one treatment or the other, and colors around the nodes indicate which treatment group that particular feature was enriched in as determined by DeSeq2 (CON=green; RPS=orange). Nodes representing 16S rRNA gene OTUs (97% similarity) are labeled with their taxonomic classification according to the SILVA database. Labels are genus names except when classification at the genus level was not possible, in which case family names are shown. Nodes representing *but* gene OTUs (97% similarity) are labeled according to the species name of their closest BLAST hit.

### Differential host response with dietary RPS

To investigate the impact of dietary RPS on immune status, and to correlate detected immune changes with bacterial community alteration, T cell populations in the cecum, ileocecal lymph node, and peripheral blood were phenotyped by flow cytometry. From this analysis we did not detect a significant difference in the abundance of CD3^+^ cells in any sample type between treatment groups, and the lack of change to the number of CD3^+^ cecal cells with dietary RPS was supported by IHC results (**Figure S3**). In the cecum, less than 1% of the CD3^+^ cells expressed the γδ T cell receptor (**Figure S2**), indicating that nearly all of the T cells in the cecum were αβ T cells.

Subsequent analysis of cecal T cell populations revealed distinct changes associated with dietary treatment. A panel of antibodies against CD3, CD4, CD8α, CD25, and FoxP3 were used to simultaneously identify 16 distinct T cell populations (**Table S5**). The relative abundance of each cell type was reported as a percent of the total CD3^+^ cells, generating a community data matrix for ecological analyses. No significant differences between groups were observed in T cell communities in the ileocecal lymph node or peripheral blood; however, a significant difference in overall cecal T cell community structure was detected between treatment groups (**Figure 7A**) (PERMANOVA p=0.001, F=12.06). Additionally, the evenness of the cecal T cell communities in CON animals was significantly reduced, meaning these communities tended to be dominated by a few cell types (**Figure 7B**, p = 0.02). Several types of cecal T cells were differentially abundant between the treatments (**Figure 7C**). More specifically, we observed an increase in several CD8α^+^ populations and a relative decrease in FoxP3^+^ cells in CON animals. These data suggest that animals fed RPS had increased abundances of T regulatory cell types associated with immune tolerance; however, these changes were limited to the cecal mucosa.

**Figure 7:**
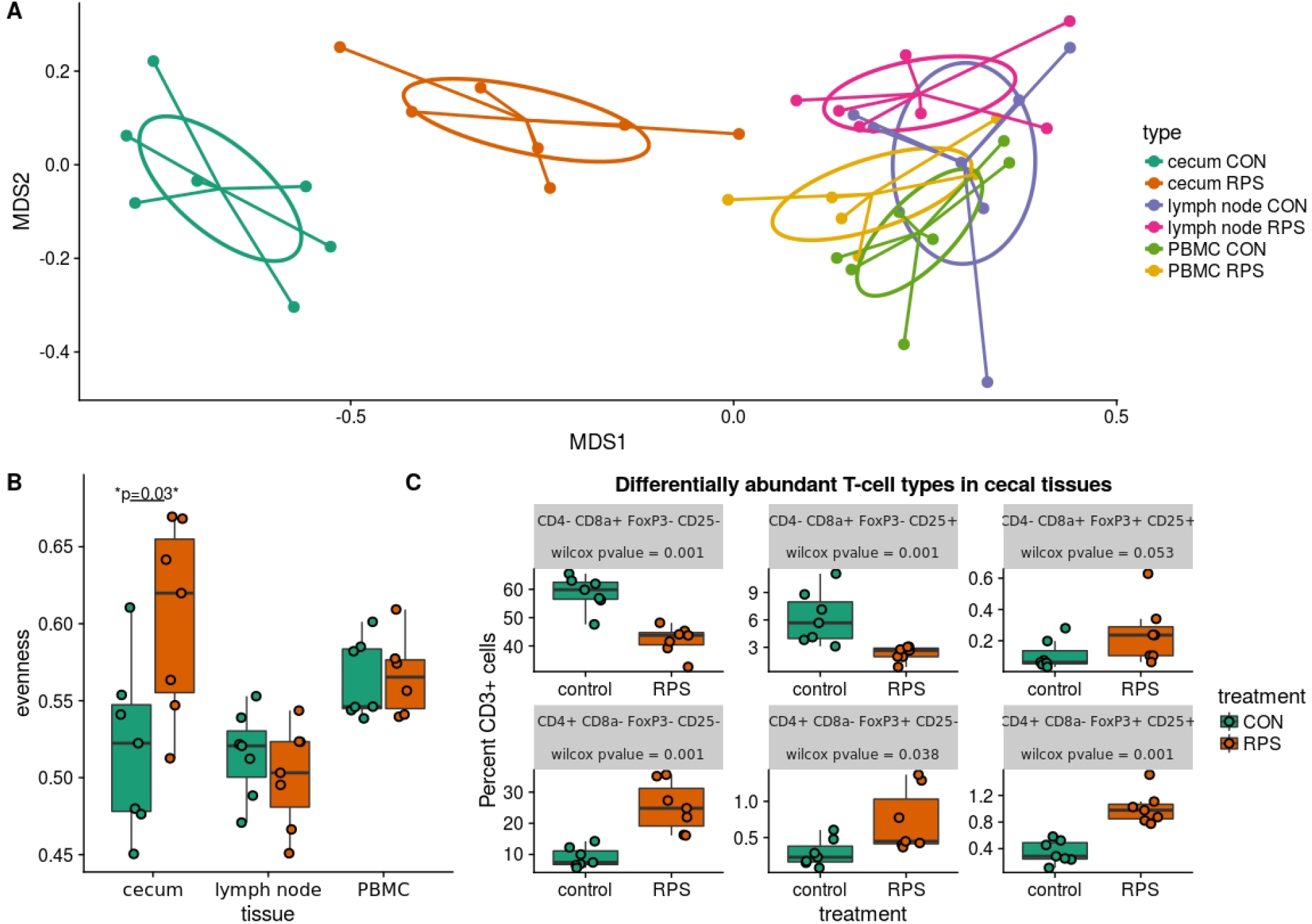
Differences in T cell populations between the RPS and CON animals at 21 days post-weaning. **A**) NMDS ordination of Bray-Curtis dissimilarities of T cell communities in various tissues. Each point represents the total T cell community for one pig in a certain tissue, and distances between points represent the similarities between communities. **B**) The evenness (Pielou’s J) of the T cell communities in various tissues. Low evenness indicates that a community is dominated by a few abundant members. **C**) Boxplots of significantly differentially abundant T cell types from the cecal T cell community.

In addition to differences in the cecal T cell populations, significant differences in the expression of genes important for barrier function were detected in the cecal mucosa. Significantly greater expression of MUC2 and IL6 was observed (p=0.03 and p=0.03), as well as a trend towards increased expression of DEF1B (p=0.16) in the RPS-fed group compared to the CON group (**Figure 8A**). Intestinal IgA is another important host-produced factor that enhances intestinal barrier function; therefore, cecal luminal contents were assayed for total IgA concentration. RPS-fed pigs trended towards increased cecal luminal IgA at necropsy (p=0.08) (**Figure 8B**). No increase in the number of IgA+ cells in cecal tissues was detected, (**Figure S3**) suggesting increased secretion of IgA from plasma cells in the cecum.

**Figure 8:**
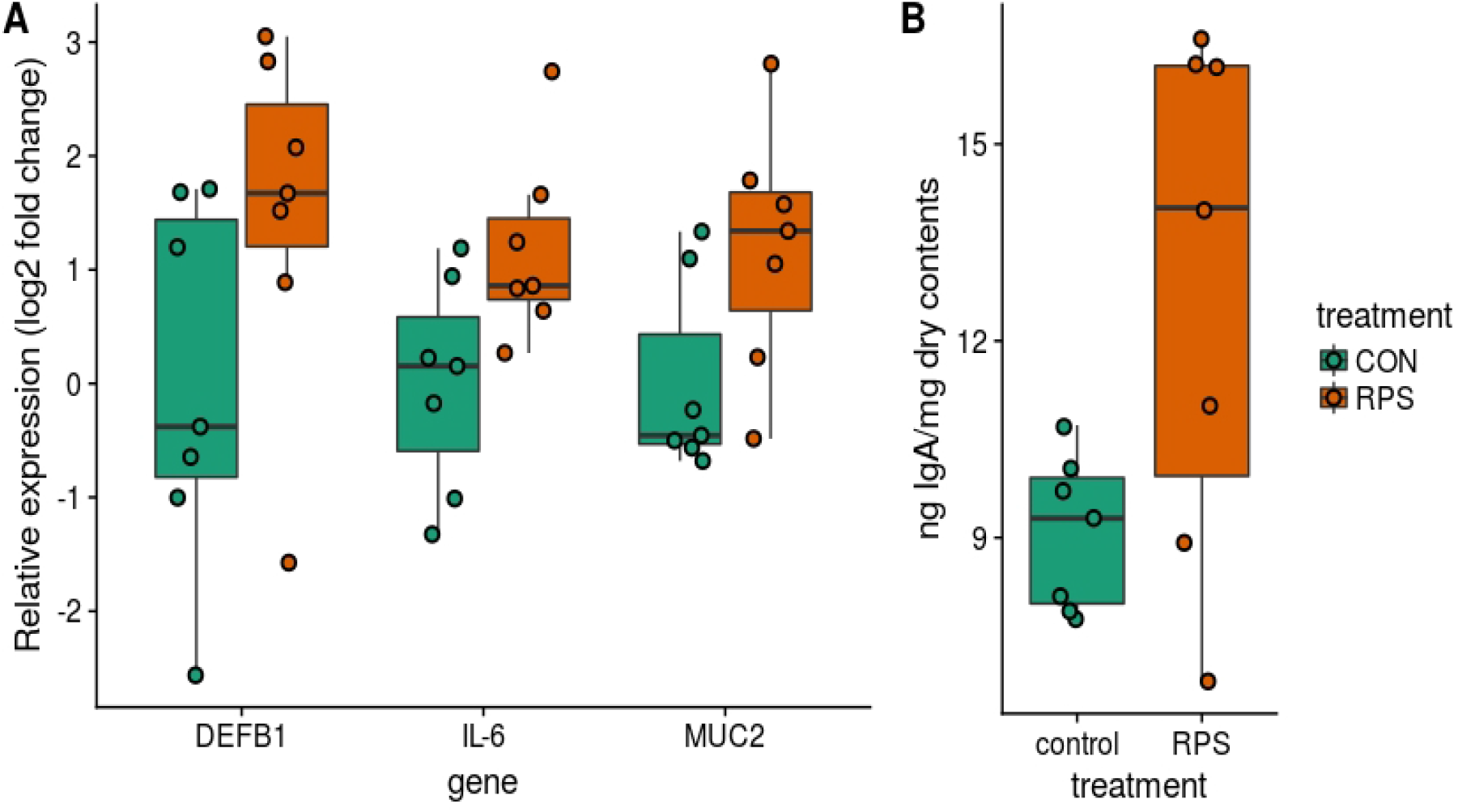
Differences in host traits associated with mucosal barrier function between CON and RPS-fed animals. **A**) mRNA expression of barrier function-associated genes from the cecal mucosa. **B**) Amount of IgA in cecal contents. Wilcoxon pvalues: DEFB1 p=0.16, IL-6 p=0.05, MUC2 p=0.05, IgA p=0.01. DEFB1= beta defensin 1, IL-6=Interleukin 6 gene, MUC2=mucin 2 gene. IgA=Immunoglobulin A.

### Bacterial members and metabolites correlated with immune status

To investigate relationships among bacterial membership, bacterial function, and host immune status in the cecum, a correlation network was constructed using the relative abundance of cecal cells [both CD3^+^ and CD3^−^ cell types were used in this analysis, (**Table S6**)], cecal 16S rRNA gene sequence diversity, cecal SCFA concentrations, cecal tissue gene expression, and luminal IgA concentrations (**Figure 9**). The results showed one discrete subnetwork associated with each respective treatment group. The subnetwork associated with pigs fed RPS (labeled subnetwork-A) was composed of features associated with immune tolerance, mucosal barrier function, and anaerobic microbial fermentation. Classic T-regulatory (CD3^+^CD4^+^CD8α^−^ CD25^+^FoxP3^+^) cells formed a central node in the RPS subnetwork along with several other CD3^+^FoxP3^+^ cell-types. Concentrations of the SCFAs butyrate, caproate, and valerate correlated with these regulatory T cells. Bacterial OTUs in the RPS subnetwork belonged to groups known for anaerobic fermentation, and several OTUs corresponded to known butyrate-producing bacterial groups, such as the genus *Megasphaera* and the family *Ruminococcaceae*. In addition, subnetwork-A contained nodes associated with an enhanced mucosal barrier: DEF1B, and IL-6 expression, and high luminal IgA concentrations.

**Figure 9:**
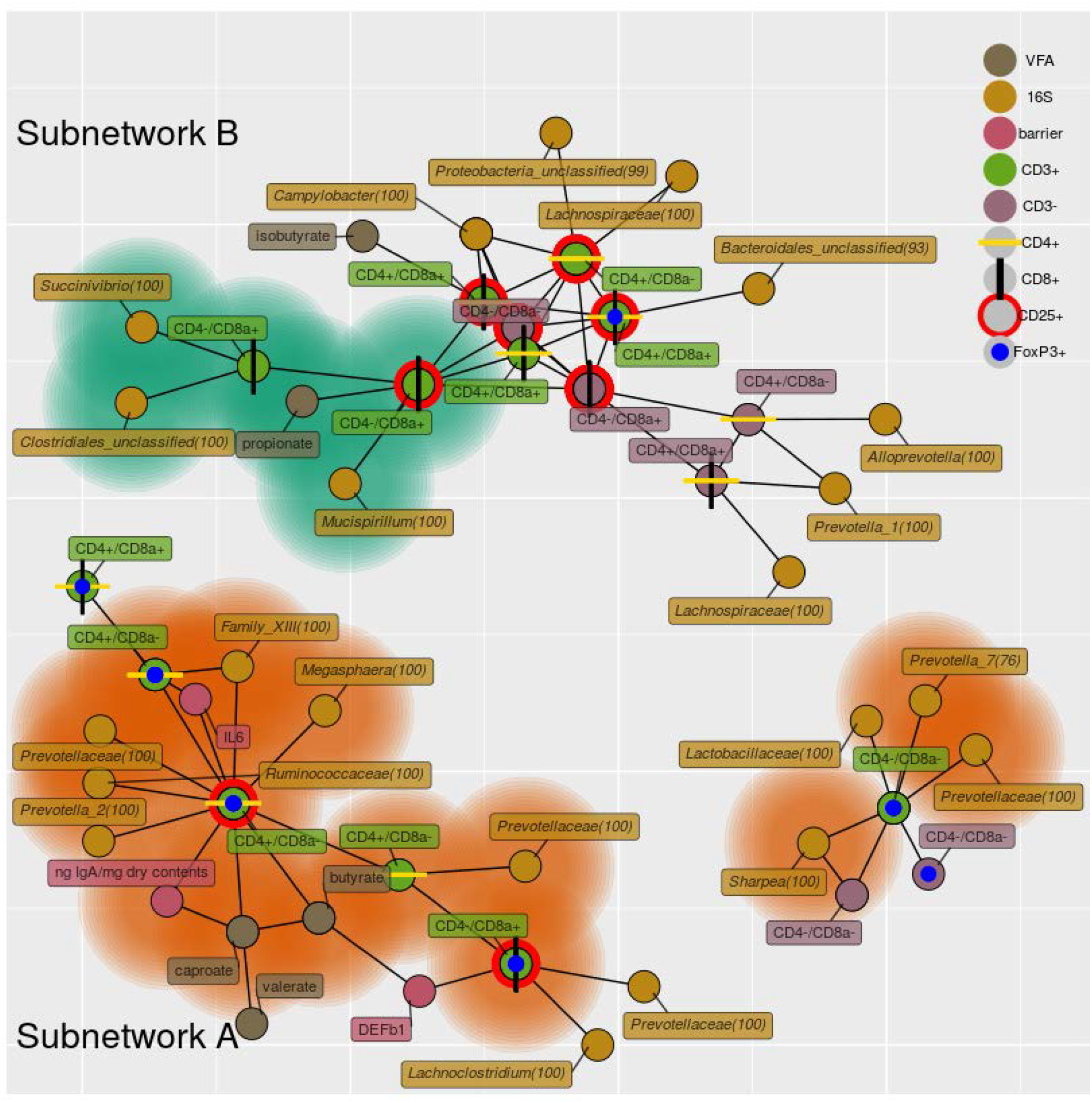
A network depicting correlations among cecal immune cells, 16S rRNA gene OTUs, cecal SCFA concentrations, cecal host mRNA expression, and cecal IgA concentrations. Only positive correlations are shown with a Spearman coefficient >0.6 and a p-value < 0.05. Bacterial nodes are OTUs labeled with their taxonomic classification according to the SILVA database. Labels are genus names except when no classification at the genus level was possible, in which case family names are shown. Nodes shades with green are features that are significantly enriched in the CON group, nodes shaded in orange are those significantly enriched in the RPS group.

The subnetwork associated with the CON diet (labeled subnetwork-B) was composed of markedly different features defined by immune activation, cytotoxic T cells, and bacteria capable of respiration. Not all of the features of subnetwork-B were significantly enriched in the CON pigs. Only cytotoxic T cells (both CD25^+^ and CD25^−^), propionate, a *Succinivibrio* OTU, and a *Mucispirillum* OTU were enriched in the CON diet. The core of subnetwork-B was composed of highly interconnected nodes, mainly cells expressing CD25 and a *Campylobacter* OTU, suggesting T cell activation and conditions conducive to microbial respiration. Several of the OTUs in subnetwork-B belong to the *Proteobacteria* phylum, members of which are known to use respiratory metabolisms (19). These results suggest that dietary RPS enhanced bacterial production of SCFAs that benefited host health by promoting epithelial barrier function, increased immune tolerance, and a reduced niche for microbial respiration relative to CON-fed pigs.

## Discussion

RPS is an accessible metabolic substrate for intestinal microbes and can lead to increased production of SCFAs, particularly butyrate, which have beneficial effects on the gastrointestinal system (20). Our results showed increased concentrations of cecal butyrate and lactate in RPS-fed animals compared to the CON animals. In the distal gut, lactate is converted to butyrate (21), and the lactate observed in the cecum was likely converted to butyrate during colonic passage as fecal lactate concentrations were very low. Supporting this idea, we observed an enrichment of butyrate–producing bacteria known to consume lactate, including *Anaerostipes hadrus* and *Megasphaera elsdenii*, in RPS-fed animals. Butyrate is well established as a bacterial metabolite of central importance for intestinal homeostasis that supports many aspects of gut health, including reducing the mucosal niche for bacterial respiration (7), increasing barrier function (4, 22), and encouraging an appropriate immune response skewing towards T cell tolerance of symbiotic microbes (23, 24).

The effects of prebiotics such as RPS are mediated through the microbiota, consequently the impact can vary greatly depending on the initial composition of intestinal bacterial communities. The large inter-individual variation present in the intestinal microbiota of mammals is well documented, swine included (25). Several studies investigating dietary resistant starch have found that prebiotic responders and non-responders can be grouped by the presence of certain microbial members (18). This phenomenon was also observed in the present study, as we detected variation in the response of individual pigs to RPS. Therefore, to optimize the effect of dietary prebiotics it may be necessary to ensure that the appropriate microbial food webs are present in the host. One study suggests that co-administration of resistant starch and a probiotic strain of *Bifidobacteria* spp. produced more desirable health outcomes than resistant starch alone (26). Our work identifies swine gut microbiota members that could be co-administered alongside RPS to potentially enhance its beneficial effects.

RPS-associated bacterial food webs were composed of organisms known to use fermentative metabolisms. Well-known fermenters such as *Bifidobacteria* spp. and *Faecalibacterium* spp. were enriched in the RPS-fed animals and are associated with intestinal health in humans (27), and other studies have shown increased abundance of these genera in pigs fed resistant starch (2, 28). Some genera enriched in the RPS-fed animals are not well characterized for their health benefits. For example, members of the genus *Clostridium sensu stricto 1* are not generally associated with positive health outcomes; however, our data suggest that members of this genus may be important for resistant starch degradation in the distal gut. Many of the differences between the treatment groups were likely a result of feedback interactions between intestinal bacterial metabolites and host tissues. Butyrate (and other SCFAs) produced by gut bacteria is oxidized by host tissues, thereby limiting the amount of oxygen available at the mucosa (7). This establishes a mucosal environment favoring microbial fermentation over respiration, and therefore more SCFA production. Increased mucosal tolerance additionally limits the release of immune-associated electron acceptors (8). In total these effects limit the niche for microbial respiration, and the types of bacteria enriched in the RPS-fed animals compared to the CON-fed animals are consistent with this model.

Increases in beneficial bacterial populations and metabolites in the RPS-fed animals were associated with positive impacts on the mucosal barrier as well as increased immune regulatory cells in the cecum. The results showed that RPS-associated microbial changes enhanced the mucosal barrier by increasing the expression of MUC2, IL-6, and DEFB1 in the cecal mucosa. IL-6 has recently been shown to be critical for maintaining the mucosal barrier (29) and has also been shown to be important for strong IgA responses (30). T cell subtypes, in particular T-regulatory cells (T-regs), and indicators of intestinal health and mucosal barrier function significantly correlated with microbial butyrate and caproate production, as well as several anaerobic *Clostridial* OTUs in the cecal mucosa of the RPS-fed animals. Butyrate is known to induce *de novo* generation and expansion of peripheral T-regs (reviewed in Zeng et. al (23)), which are critical for intestinal homeostasis and gut barrier function (31). T-regs moderate immune responses to commensal microbes, reducing intestinal inflammation and subsequent mucosal availability of immune-derived electron acceptors, thereby limiting microbial respiration (7). T-regs can promote mucosal IgA responses, helping the host exert control over its microbial partners (23, 32, 33), and cecal luminal IgA levels were increased in RPS-fed pigs and correlated with the abundance CD4^+^ T-regulatory cells. CD8α^+^ T-regulatory cells (CD4^−^ CD8α^+^ CD25^+^ FoxP3^+^) are less well studied than CD4^+^ T-regs, but recent work indicates they are an important regulatory cell type in humans and mice (34), and we detected these cells in the cecum of pigs. Though their direct role in intestinal health is unclear, the network analysis suggests that animals with greater relative abundance of CD8^+^ T-regs correlated with greater expression of DEFB1.

Though they were healthy, the CON group exhibited markers of reduced immune tolerance and greater abundances of potentially invasive bacteria in their mucosal tissues. Specifically, CON-fed animals had more cytotoxic T cells (CD8α^+^ cell types) compared to RPS-fed pigs, although they were unlikely active given the lack of pathological changes in the cecum. Similarly, bacteria enriched in the CON animals have previously been associated with intestinal inflammation, dysbiosis in humans and mice (35), and utilize respiration as their preferred metabolism. In particular, members of the genus *Mucispirillum* have been shown to thrive in inflamed, oxygenated mucosal environments (16). Our results showing correlations between the abundance of *Mucispirillum* and cytotoxic T cells suggest that an immune cell activity may play a role in expanding the niche for this organism in the CON pigs. Similarly, we detected enrichment of *Helicobacter* 16S rRNA in the cecum and feces from CON-fed animals. Bacteria from this genus can be facultative intracellular pathogens (36), and it has been proposed that *non-H. pylori Helicobacter* species may be a cause of irritable bowel disease (IBD) in humans (29). These observations suggest that though the CON pigs lacked distinct pathology, the mucosae of these animals were more amenable to colonization by potentially pathogenic organisms that utilize respiratory metabolisms, relative to the RPS pigs.

Local intestinal inflammatory responses have previously been shown to occur in swine early in weaning (37), and though no overt inflammation was observed in these tissues in the current study, we detected immune cell types associated with recent immune activity in both treatment groups. In the cecal tissue network analysis, many nodes residing in subnetwork B were equally represented in both the CON and RPS groups. For example, double-positive (CD4^+^CD8α^+^) CD25^+^ T cells were prominent members of subnetwork B and were not differentially abundant between treatments; these cells have been shown to be common in swine and represent activated, memory effector cells (38). Interestingly, a *Campylobacter* OTU, was correlated with the abundance of these cells and several other CD25^+^ cell types. This observation suggests that certain bacterial groups, particularly *Proteobacteria*, may benefit from the local environmental changes associated with recent immune activity, such as the release of reactive nitrogen or oxygen species. While the memory cells may not be the source of reactive nitrogen or oxygen species, their presence indicates prior perturbation of the mucosal site. The idea that *Proteobacteria* thrive using products of the immune response is well established for *Salmonella* in mice (8, 9) and our data suggest that this model warrants further investigation for other bacterial species and hosts.

## Conclusions

This study demonstrated that dietary intake of RPS had beneficial impacts on the intestinal health status of weaning pigs, including increased markers of mucosal barrier function, immune tolerance, and increased abundances of potentially beneficial bacterial populations. Our study revealed the specific bacterial interactions involved in the bacterial breakdown of resistant potato starch, and identified specific bacterial groups that could be co-administered with RPS to enhance its effects. Additionally, this work revealed specific correlated changes between the commensal microbiota and the mucosal immune system that can be used to inform future strategies to modulate the microbiota to support health. These data provide valuable insights into the host-microbe interactions in the intestinal mucosa of swine, furthering our knowledge of the mammalian hindgut ecosystem. Finally, pigs are recognized as a relevant translational model for human health, and research to enhance intestinal health in pigs provides insights for enhancing human health as well.

## Materials and Methods

### Experimental design

Ten pregnant, Large White crossbred sows were delivered two weeks prior to farrowing, and farrowed onsite. At 14 days-of-age, piglets were offered non-amended Phase 1 starter diet (**Table S1**). At 21 days-of-age, piglets were weaned, and separated into two treatment groups. Treatment groups consisted of two pens of seven piglets for a total of 14 piglets in each treatment group, each group had equal representation from all litters. The control group (CON) continued to receive non-amended Phase 1 Starter Diet. The treatment group was fed Phase 1 Starter Diet amended with 5% raw potato starch (RPS; MSP Starch Products Inc., Carberry, Manitoba, Canada). At 12 days post-weaning (33 days-of-age), the CON group was switched to non-amended Phase 2 Diet and the RPS group switched to Phase 2 Diet amended with 5% raw potato starch (**Table S1**). At 21 days post-weaning (42 days-of-age), seven piglets from each group (3 from one pen and 4 from a second pen) were humanely euthanized by injection of sodium pentobarbital (Vortech Pharmaceuticals). All animal procedures were performed in compliance with the National Animal Disease Center Animal Care and Use Committee guidelines and review.

### Sample Collection

Piglets were weighed at weaning and necropsy. Fecal samples were collected at 0, 12, 15, 19, and 21 days post-weaning. Feces were collected fresh and transported on ice, aliquoted for downstream applications, and stored at −80 °C. Prior to euthanasia, peripheral blood was collected into sodium citrate cell-preparation tubes (BD Pharmingen). At necropsy, cecal contents were collected into RNALater and stored at 4 °C until RNA extraction, and snap frozen and stored at −80 °C. Sections of cecal tissue were gently rinsed with phosphate-buffered saline (PBS) and the mucosae were scraped with a sterile cell lifter. One portion of mucosal scrapings was immediately stored in RNALater at 4 °C for host RNA extraction and another frozen at −80 °C for bacterial DNA extraction. Ileocecal lymph node and sections of cecal tissue were collected in the appropriate buffer on ice and immediately processed for flow-cytometry.

### Immunohistochemical Staining (IHC)

Fresh cecal tissues were formalin-fixed and sections prepared using standard histological techniques. Details for CD3 and IgA staining are available in the supplement. Slides were scanned into Spectrum Version 11.2.0.780 (Aperio Technologies, Inc.) and Aperio ImageScope was used for annotation and to quantify cell populations. Cell counts were obtained using a nuclear algorithm on Aperio ScanScope software and are reported as cells/mm^2^.

### Phenotypic analysis by flow cytometry

At necropsy, ~2 g of gently rinsed cecal tissue was placed in complete RPMI (RPMI 1640 [Life Technologies; Grand Island, NY] supplemented with 10% fetal calf serum [FCS, Omega Scientific; Tarzana, CA], L-glutamine [Life Technologies], 25 mM HEPES [Sigma; St. Louis, MO], essential amino acids and antibiotics [Sigma]) and stored on ice until processing. A previously described protocol was adapted for isolation of both epithelial cells and lamina propria cells from cecal tissue (39), with some modifications (Supplementary Information). Cells from peripheral blood and ileocecal lymph nodes were isolated as previously described (40). Approximately 10^6^ cells per tissue were used for flow cytometric analysis.

Cells were stained with Zombie Yellow Viability dye (Biolegend, San Diego, CA), followed by incubation with fluorescently-conjugated anti-porcine monoclonal antibodies (BD Biosciences, San Jose, CA (except as noted)). Antibodies used included anti-porcine CD3 (clone BB23-8E6-8C6), CD4 (clone 74-12-4), CD8α (clone 76-2-11), CD25 (clone K231.3B2, Southern Biotech), γδTCR (clone MAC320), and FOXP3 (clone FJK16s). For staining of FoxP3 the Intracellular Nuclear Staining Kit (Biolegend) was used. Data were acquired on a BD LSRII instrument and analyzed with FlowJo Software (FlowJo LLC, Ashland, Oregon). Representative flow plots and gating strategy are available (**Figures S1 and S2**).

### RNA extraction, cDNA synthesis, and RT-qPCR of cecal tissue

Host RNA was extracted using the TriReagent (Life Technologies)-modified protocol with the PowerLyzer UltraClean Tissue & Cells RNA Isolation Kit (MoBio Laboratories, Inc.). Homogenization in TriReagent was carried out in a Thermo Savant FastPrep^®^ FP120 Cell Disrupter (Qbiogene, Inc., Carlsbad, CA). An on-column DNAse step was included (On-Spin Column DNase I Kit, Mo Bio Laboratories, Inc.). QuantiTect Reverse Transcription Kit (Qiagen, Valencia, CA) was used for cDNA synthesis.

Gene expression was measured using the TaqMan^®^ Universal Master Mix II system (Applied Biosystems, Foster City, CA). Cycling conditions were 40 cycles of 95°C for 15 sec and 60°C for 1 minute. The gene β-actin was used to normalize the expression of target genes according to the 2^-ΔΔCq^ method (41). Primers and probes are described in **Table S2**.

### Microbial community analysis

Microbial nucleic acids were extracted from feces and cecal contents using the PowerMag fecal DNA/RNA extraction kit (MoBio). Only cecal contents were used for RNA isolation. RNA samples were treated with DNase Max (MoBio) kit, and converted to cDNA using the High Capacity cDNA Synthesis Kit (Applied Biosystems). Amplicons of the V4 region were generated and sequenced in accordance with the protocol from Kozich et al. (42). Amplicons of the butyryl-CoA:acetate CoA transferase (*but*) gene were generated and sequenced using the protocol from Trachsel et al., 2016 (43). Normalized pools for the 16S rRNA gene library and *but* gene libraries were sequenced on a MiSeq (Illumina) using 2×250 V2 and 2×300 V3 chemistry, respectively.

### IgA measurements

Cecal contents (~250 mg) were lyophilized, resuspended in extraction buffer (10mM Tris, 100mM NaCl, 1mM CaCl2, 0.5% Tween-20, 1 tablet cOmplete™, EDTA-free Protease Inhibitor Cocktail (Roche, Branford, CT) per 100 mL) at 1mL extraction buffer per 30 mg freeze-dried cecal contents, and vortexed on high for 10 minutes. Debris was pelleted by centrifugation at 5000 x g and dilutions of the supernatant were used in the Pig IgA ELISA Quantitation kit (Bethyl Laboratoties, Montgomery, TX). Results are reported as ng IgA/mg dry contents.

### Short-chain fatty acid (SCFA) measurements

One gram of material (cecal contents or feces) was suspended in 2 mL PBS, vortexed for one minute, and debris was pelleted by centrifugation at 5000 x g for 10 minutes. Supernatant (1 mL) was added to heptanoic acid internal standards. Butylated fatty acid esters were generated as described (44), and analyzed using an Agilent 7890 GC (Agilent, Santa Clara, CA).

### Data analysis

Unless otherwise stated, Wilcoxon tests were used to test for statistical differences. Both the 16S rRNA gene and *but* gene amplicon data were clustered into operational taxonomic units (OTUs) with 97% similarity in mothur using the Miseq SOP (45). 16S rRNA gene sequences were aligned to the SILVA reference alignment, and *but* sequences were aligned to an alignment of *but* reference sequences downloaded from RDP’s fungene database (46). Error rates calculated by sequencing mock communities (47) for 16S rRNA genes and *but* genes were 1.3e-06 and 2.2e-03 errors per basecall, respectively.

The R package vegan (48) was used to carry out ecological analyses. Community structure similarity analyses were performed by calculating Bray-Curtis dissimilarities, and statistical testing was accomplished using vegan’s adonis and betadisper functions. Differential abundance was determined using the DeSeq2 package (49). Correlations for network analysis were calculated with CCREPE (50) for compositional data or the rcorr function from the hmsic R package. Network layout was performed with the geomnet R package (30). Only significant, positive correlations with a Spearman coefficient of at least 0.6 are shown.

### Data availability

Raw sequence data is available in the SRA under project accession number PRJNA476557. All processed data and R scripts used in this analysis are available at https://github.com/Jtrachsel/RPS-2017.

## Acknowledgements

We are grateful to the NADC animal care staff for their efforts. We thank Jennifer Jones, Zahra Olson, Nicole Hasstedt, and Sam Humphrey for technical assistance, and Haley Jeppson for help with R coding and figure consultation. Mention of trade names or commercial products in this publication is solely for the purpose of providing specific information and does not imply recommendation or endorsement by the U.S Department of Agriculture. USDA is an equal opportunity provider and employer.

## Author Contributions

JT, CL, and HKA conceived of and planned the study. NG formulated and prepared the diets, and provided guidance on diet phases. CL and CB collected the gene expression, flow cytometry, IgA, and immunohistochemistry data. JT collected microbial community data and SCFA data. JT performed the data analysis. JT, CB, HKA, and CL wrote and revised the manuscript.

